# Unraveling the archaeal virosphere: diversity, functional and virus-host interactions

**DOI:** 10.1101/2025.11.27.691053

**Authors:** Yang Zhao, Pengfei Liu, Meiling Feng, Rong Wen, Zhihao Zhang, Xuefeng Zhang, Xingyu Huang, Hongfei Chi

## Abstract

Archaea, the third domain of life, play critical roles in global biogeochemical cycles. However, their virosphere, particularly the proviruses which integrated into host genomes, remains largely unexplored. To systematically reveal the landscape of archaeal proviruses, we conducted large-scale mining of public and in-house genomic datasets spanning all presently known archaeal phyla. We identified 9,697 archaeal proviruses across 19 archaeal phyla and 366 families, which clustered into 9,123 viral operational taxonomic units (vOTUs). Among these, 97.2% represent novel viruses, and 81.3% could not be classified at the family level, substantially expanding the known diversity of archaeal viruses. Host range analysis revealed that many proviruses exhibit broad infectivity across archaeal lineages, with some even capable of cross-domain infection. Genomic analysis identified 178 distinct types of antiviral systems in archaeal hosts, encompassing multiple CRISPR-Cas variants and restriction-modification (RM) systems. Meanwhile, we detected 747 anti-defense genes encoded by 710 proviruses, such as anti-CRISPR and anti-RM, directly corroborating the ongoing evolutionary arms race between archaeal hosts and their viruses. Additionally, we identified 532 auxiliary metabolic genes (AMGs) within archaeal proviruses that are involved in key processes including carbon, nitrogen, and sulfur metabolism, indicating their potential to reprogram host metabolic pathways and thereby influence biogeochemical cycling. This study establishes a systematic global genomic atlas of archaeal proviruses, advancing our understanding of their distribution and diversity while laying the groundwork for future investigations into how AMG-mediated processes influence archaeal metabolism and ecosystem functions.

## INTRODUCTION

Archaea, the third domain of life on Earth, represent a major group of single-celled microorganisms alongside Bacteria and Eukaryotes^1, 2^. The taxonomic classification of Archaea has been continuously revised and expanded. Initially, all archaea were classified into two primary phyla—Euryarchaeota and Crenarchaeota^1, 3, 4^. Over the past three decades, advances in phylogenetic and genomic analysis have propelled archaea into the research spotlight, spurring growing interest and comprehensive exploration across diverse habitats, from extreme to conventional environments. Currently, the SILVA database, based on small ribosomal subunit gene sequencing, lists approximately 30 archaeal phyla encompassing about 20,000 species^5^. In contrast, the genome-based classification system of the Genome Taxonomy Database (GTDB) has 20 phyla with 6,968 species^6–8^. Within the increased discovery of new species, the archaeal taxa are classified into three superphyla: TACK, DPANN, and Asgard^9–11^. The TACK and DPANN superphyla were originally named based on their constituent phyla but have since been reclassified and expanded. For example, the DPANN superphylum initially included Diapherotrites, Parvarchaeota, Aenigmarchaeota, Nanohaloarchaeota, and Nanoarchaeota, but now comprises more than ten phyla, such as Aenigmatarchaeota, Altiarchaeota, Huberarchaeota, Iainarchaeota, Micrarchaeota, Nanobdellota, Nanohaloarchaeota, Undinarchaeota, along with other tentative groups^6^. These phyla of microorganisms mostly hard to be cultured and completely sequenced, and they are often referred to as “microbial dark matter”^11^. Nevertheless, genomic data from uncultivated archaea have accumulated exponentially in recent decades^9, 11, 12^, revealing their crucial ecological roles in diverse habitats^13–15^.

In parallel, the rapid advancement of high-throughput sequencing technologies and automated bioinformatic workflows has significantly accelerated the discovery of archaeal viruses. To date, most isolated archaeal viruses have been obtained from hyperthermophilic or hyperhalophilic hosts within the phyla Thermoproteota and Euryarchaeota, with only a limited number described to infect methanogenic archaea^16^. Recent metagenomic studies are continually expanding this view, revealing a broad spectrum of novel, uncultivated archaeal (pro)viruses that have now been formally classified into established or new viral families^17–20^. Archaeal proviruses are defined as viral genomes that have integrated into the genomic DNA of the host cell^21^. The majority of known archaeal proviruses include head-tailed viruses and pleolipoviruses, the latter being a group of pseudo-spherical and pleomorphic archaeal viruses composed of a membrane vesicle enclosing a DNA genome^21, 22^. These are predominantly identified in hosts from the Methanobacteriota, Thermoproteota, and Halobacteriota phyla^23–25^.

Over evolutionary time, some proviruses are immobilized by their host and the sequences encoding beneficial traits conserved in cells through natural selection, thereby profoundly influencing host evolution and metabolism^25, 26^. One way this virus–host interaction manifests is through viral auxiliary metabolic genes (AMGs), which can enhance or redirect host metabolic processes to facilitate viral replication^27^. For instance, AMGs identified in marine Thaumarchaeota viruses encode a key subunit (*amoC*) of the ammonia monooxygenase, a central enzyme in nitrification, potentially enhancing the host’s energy metabolism and viral replication during infection^28^. Conversely, persistent viral infection has driven the evolution of sophisticated host defense mechanisms. Among these, the CRISPR-Cas system stands out as a highly adaptive immune system that archives fragments of past viral infections to confer sequence-specific immunity upon re-infection^29, 30^. These archived CRISPR spacer sequences also constitute a valuable resource for discovering novel viruses and elucidating virus–host dynamics^31^. Although recent studies have begun to characterize CRISPR-Cas diversity in specific lineages such as the DPANN superphylum, Asgardarchaeota, and *Bathyarchaeia*^18, 20, 32^, a comprehensive assessment of CRISPR and other defense systems—including classical mechanisms such as restriction-modification—across the broader archaeal domain remains lacking.

To address the knowledge gaps in archaea provirus diversity, classification, and virus-host interactions, we conducted systematic genomic mining of both public repositories and in-house datasets. This study aimed to: (i) characterize the diversity, classification, and host range of archaeal proviruses; (ii) identify antiviral systems in archaeal hosts and associated anti-defense mechanisms encoded by proviruses; and (iii) annotate AMGs carried by archaeal proviruses and assess their potential metabolic roles. Our analyses reveal a vast and phylogenetically diverse array of archaeal proviruses, which encode a variety of anti-defense system and AMGs. These findings significantly expand the known diversity of archaeal viruses and provide new insights into virus–host evolutionary dynamics and functional interactions in archaea.

## RESULTS AND DISCUSSIONS

### The overall landscape of provirus elements in archaea

To investigate provirus associated with archaea, we analyzed 21,989 archaeal genomes (completeness ≥ 50% and contamination rate < 10%) from both public databases (98.2%, *n*=21,593) and our own dataset (1.8%, *n* = 396) using two virus identification methods (Methods; Table S1). A total of 9,697 provirus regions (≥ 5 kb) were identified in 19.1% (4,196/21,989) of these genomes, spanning 60.7% (366/603) of archaeal families represented in the dataset (Figure 1a). The number of identified proviruses varied greatly among the strains, with approximately 55.0% (2,308) of strains harboring only one provirus, while 90 strains carried more than 10 proviruses (Figure 1b). Notably, over 50 proviruses were detected in two archaeal genomes (GCA_022759085.1 and QYJanS1_maxbin.027), which were metagenome-assembled genomes (MAGs). The huge variation in the number of proviruses indicates a highly uneven distribution of proviruses among archaea.

**Figure 1.**
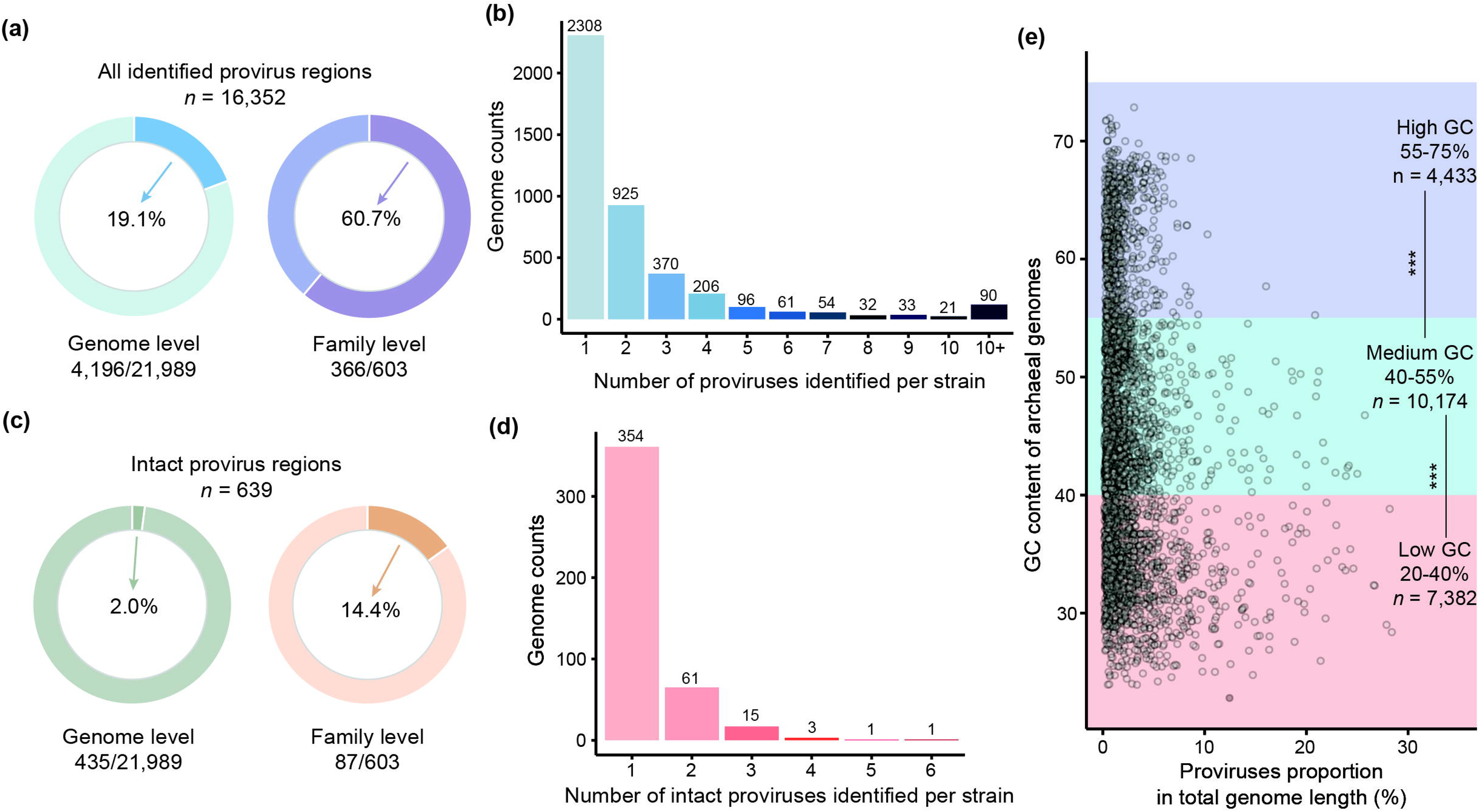
The overall landscape of archaeal proviruses. (a) The detection rates of all identified provirus regions at the genome level and the family level. (b) The number of provirus regions carried by each strain. (c) The detection rates of intact proviruses at the genome level and the family level. (d) The number of intact proviruses carried by each strain. (e) The proportion of provirus length relative to host genome size. The archaeal genomes are divided into three groups based on their GC contents: the “low GC” group: archaeal genomes with GC content between 20-40%; the “Median GC” group: archaeal genomes with GC content between 40-55%; the “high GC” group: archaeal genomes with GC content between 55-75%. Significance tests were performed using the nonparametric Mann-Whitney U test, and the two-tailed p values were calculated. \*\*\**p* < 0.001.

Most of the predicted provirus regions are incomplete provirus fragments, while 544 provirus regions are marked as intact proviruses (evaluated as “complete/high quality” by CheckV), distributed in 435 (2.0%) genomes and 87 (14.4%) archaeal families (Figure 1c). Of the 435 archaea genomes that harbor intact proviruses, the vast majority (81.4%) carried less than 2 intact proviruses (Figure 1d). However, several archaeal hosts are parasitized by multiple intact proviruses, especially in *Ignisphaera* sp. isolate 2016_B02_sed_C_13, of which as many as six intact provirus genomes were identified. Life strategy predictions further revealed that the proportion of virulent proviruses (5,852; 60.3%) was markedly higher than that of temperate ones (2,399; 24.7%) (Table S2).

Previous studies have reported that the number of proviruses is positively correlated with the size of their host genomes^33^. Here, our analysis revealed a significant, albeit weak, positive correlation (Spearman *ρ* = 0.221, *p* = 2.8^e-241^; Figure S1a) between host genome size and the number of identified proviruses. Moreover, we observed a similarly significant but weaker positive correlation between host genome size and provirus genome proportion, which represents the proportion of the total host genome length occupied by proviruses (Spearman *ρ* = 0.209, *p* = 1.4^e-214^; Figure S1b). This weak correlation suggests that host genome size has a limited impact on provirus carrying frequency and genomic proportion, which might also be influenced by other genomic characteristics. Given the established impact of Guanine and Cytosine (G+C) content on species ecology, distribution, adaptation, and lifestyle^34, 35^, we investigated its association with provirus carriage in archaea. The 21,989 archaea were divided into three groups: “low GC” (20-40%, *n* = 7,382), “medium GC” (40-55%, *n* = 10,174), and “high GC” (55-75%, *n* = 4,433) groups. Subsequently, a comparison was made regarding the differences in provirus genomic proportion and the number of proviruses among these groups. Clearly, archaea with high GC content possess a significantly lower provirus genomic proportion (*p* < 0.001; Figure 1e) and a smaller number of proviruses within their genomes (*p* < 0.001; Figure S1c), whereas archaea with low GC content exhibit the opposite trend.

We further examined the distribution of proviruses across archaeal families. First, the majority of families (86.9%, *n* = 525) had provirus regions detected in fewer than half of their strains (Table S3). Only 30 families (5.0%) showed proviruses in all their strains (e.g., B15-G16, JALHTG01, JAEHCK01). However, each of these 30 families was represented by only 1-2 genomes in our genome dataset. In addition, *Methanobacteriaceae* had the most provirus regions (1,075), followed by *Bathycorpusculaceae* (376), *Nitrosopumilaceae* (350), and GW2011-AR-1 (342) (Table S3). This distribution pattern may be related to the significant imbalance of archaeal genomes in the dataset, as most genomes belong to the *Nitrosopumilaceae* (1,078), *Methanobacteriaceae* (878), GW2011-AR-1 (585), and *Bathycorpusculaceae* (410). To take into account this possible bias, we compared the number of proviruses from 10 randomly selected genomes (Methods) and performed this step by bootstrapping (100 times) to sample all the genomes in our database. We found that three families, *Caldarchaeaceae*, *Calditenuaceae*, and *Methanoperedenaceae*, were richer in proviruses (Figure S2). Among these, *Methanoperedenaceae* is broadly distributed across diverse habitats (e.g., groundwater, bioreactors, lakes, and soil), whereas *Caldarchaeaceae* and *Calditenuaceae* are predominantly isolated from hot springs (Table S4). At the genus level, significant provirus enrichment was observed in *Caldarchaeum*, *Methanoperedens*, HRBIN02 of *Calditenuaceae*, *Bathycorpusculum*, *Methanobrevibacter*, *Nitrosocaldus*, and *Sulfolobus* (Figure S2). In contrast, no significant enrichment was detected at higher taxonomic ranks (phylum, class, or order), indicating this phenomenon is confined to the family and genus levels (Figure S2).

### Diversity, taxonomy and host range of archaeal proviruses

To explore the diversity of archaeal proviruses, we performed clustering analysis on 9,697 identified proviruses using thresholds of ≥ 95% average nucleotide identity and ≥ 85% alignment fraction (Methods), yielding 9,123 viral operational taxonomic units (vOTUs). Overall, 6,927 vOTUs (75.9%) were classified to known taxa using the Lowest Common Ancestor method (Methods; Figure 2; Table S5). These proviruses spanned five DNA virus realms: *Duplodnaviria* (72.4%, *n* = 6,604), *Varidnaviria* (2.0%, 79), *Adnaviria* (0.9%, 86), *Monodnaviria* (0.4%, 38), and *Singelaviria* (0.2%, 20) (Figure 2a; Table S5). At class level, the majority (72.4%) of vOTUs were assigned to the class of *Caudoviricetes* within the phylum *Uroviricota* (Figure 2a), which comprises head-tailed viruses ubiquitous in natural environments and human hosts^36, 37^. Only 1,707 (18.7%) vOTUs could be resolved to the family level, indicating high genetic diversity among archaeal proviruses and the presence of numerous previously undetected novel viruses. These vOTUs were distributed in 87 different families, mainly including *Kyanoviridae* (386, 4.2%), *Herelleviridae* (75, 0.8%), *Autographivirales* (71, 0.8%), *Straboviridae* (71, 0.8%) and *Hafunaviridae* (71, 0.8%) (Figure 2a).

**Figure 2.**
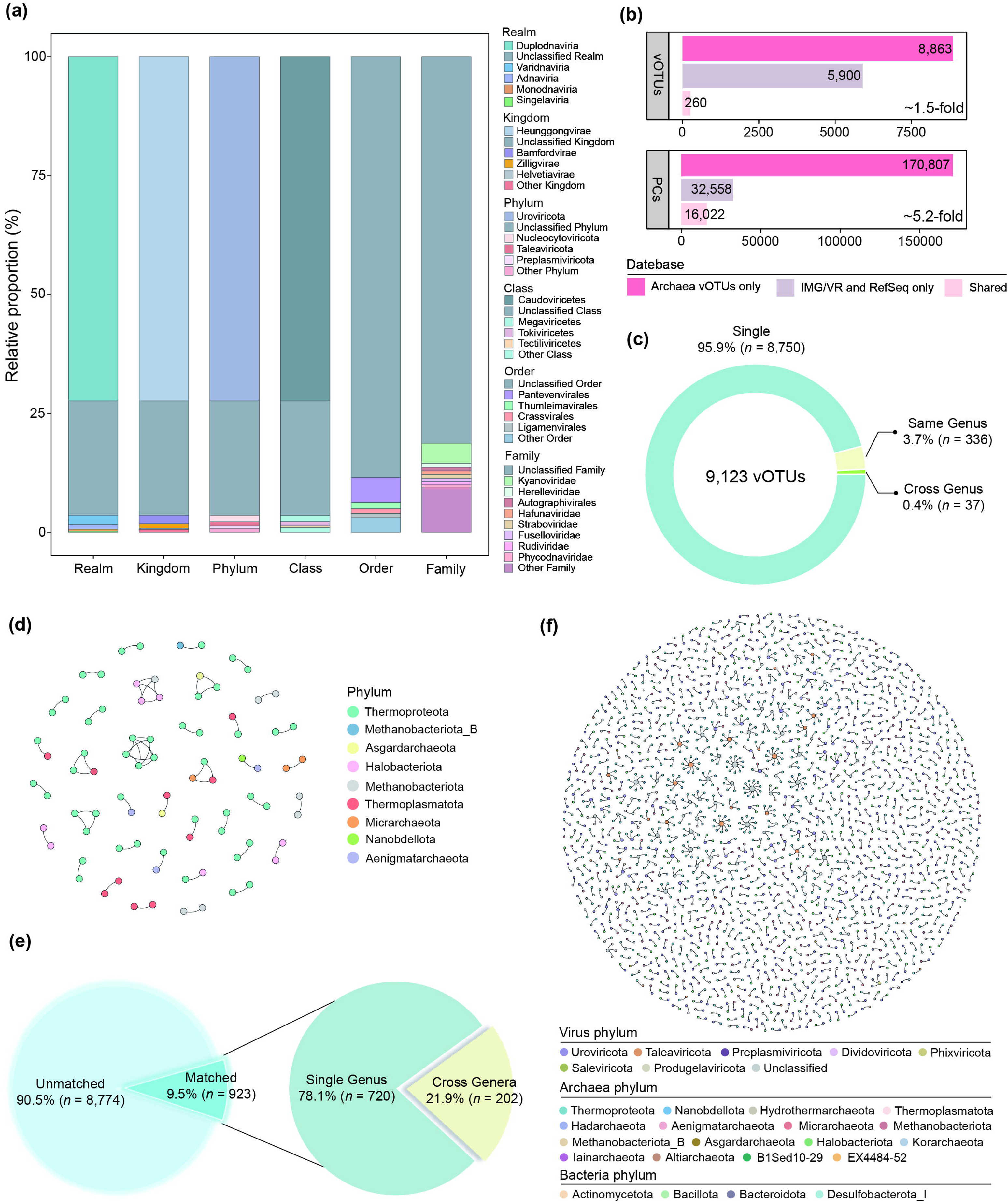
Diversity, host range, and host-provirus interactions of the archaeal proviruses. (a) The proportion of archaeal provirus communities at the phylum, class, order, and family levels. (b) Number of vOTUs and their PCs in the archaeal provirus dataset, the IMG/VR v4 and NCBI Viral RefSeq, and shared by the databases. (c) 9,123 vOTUs were classified into three categories based on their clustered host range. Single: this vOTU is only integrated into a single archaeal genome; same genus: This vOTU may infect different genomes of the same archaeal genus; Cross genus: This vOTU may infect genomes belonging to different archaeal genus. (d) Integrating associations between host phyla for each cross-genus clustered vOTU. (e) The CRISPR spacer sequence targeting situation of archaeal proviruses. The left chart shows that the vast majority (90.5%) of proviruses were not targeted by any spacers. The right chart details the host range of the targeted provirus. (f) This network describes the relationship between the provirus and its host based on CRISPR spacer predictions.

In addition, to assess the novelty of the archaeal proviruses in this study, archaeal virus contigs (≥5 kb) and their proteins were extracted from NCBI RefSeq and IMG/VR for comparative analysis (Methods). The results demonstrated that our provirus dataset significantly expands the known diversity of archaeal viruses, with a 1.5-fold increase at the vOTU level and a 5.2-fold increase at the protein cluster (PC) level (Figure 2b), implying archaea as a large reservoir of unknown viruses. Notably, although host classification was consistent between most archaeal viruses in public databases and the archaeal proviruses they clustered with, eight instances of mismatched host classification suggest potential cross-species provirus transmission. Supporting this, analysis of vOTU host distribution revealed that 99.6% (*n* = 9,086) of the 9,123 vOTUs were restricted to a single archaeal genus, whereas 0.4% (*n* = 37) spanned different archaeal genera (Figure 2c). Most of these broadly distributed vOTUs were confined to two genera within the same phylum (Figure 2d). Whereas 13 vOTUs crossed phylum boundaries. Among them, two harbored integrase encoding genes, including unclassified vOTU GCA_038894755.1_scaffold_25, which is predicted to potentially infect hosts within the phyla Thermoproteota and Aenigmatarchaeota, and *Caudoviricetes* vOTU GCA_038873995.1_scaffold_12 exhibited a predicted host range spanning Thermoproteota, Thermoplasmatota, and Micrarchaeota. These findings indicate that certain archaeal proviruses possess an unusually broad integration capacity, potentially facilitating horizontal gene transfer across highly divergent archaeal lineages.

The presence of spacers in CRISPR systems allows us to trace the infecting history of archaeal proviruses. Using the golden standard CRISPR spacer targeting method, we found that 90.5% (8,775/9,697) of archaeal proviruses did not match any spacers (Figure 2e). Among the 922 proviruses that were targeted by spacers, 78.1% (*n* = 720) were targeted exclusively by spacers from a single genus, while the remaining 21.9% (*n* = 202) exhibited a broad host range, infecting between 2 and 11 distinct genera (Figure 2e). Among these, the unclassified provirus GCA_038838075.1_scaffold_44 (11) and GCA_038845585.1_scaffold_38 (10) infected the highest number of genera. Furthermore, 296 of the spacer-targeted proviruses showed no targeting by spacers derived from their own isolation genus (Figure 2f). These results suggest that the vast majority of archaeal proviruses (93.5%, 9,071/9,697) may be “Silencers”—proviruses capable of infecting archaeal hosts without leaving an immune record in the host’s defense system^38^. Notably, 37 archaeal proviruses were found to be capable of infecting bacteria. For instance, the *Phixviricota* provirus GCA_030623225.1_k141_277094 infected three genera within the Bacteroidota phylum (*Bacteroides*, *Phocaeicola*, and *Phocaeicola_A*), while the unclassified provirus GCA_949320185.1_KLAP88_365180 infected three genera across three different bacterial phyla: *Collinsella* (Actinobacteriota), *Desulfocurvus* (Desulfobacterota_I), and *Absicoccus* (Bacillota) (Figure 2f).

### Antiviral systems in archaea

Archaea have evolved diverse and complex antiviral systems to protect against viral infection^39^. Among these, the CRISPR-Cas system is the most common antiviral defense strategy, providing adaptive immunity to the host by memorization and the recognition of past viral attacks^30^. A total of 1,297 complete CRISPR-Cas systems were identified in 4.9% (1,070/21,989) of archaeal genomes analyzed, and these CRISPR-Cas systems can be categorized into 2 classes, 5 types, and 29 subtypes (Figure S3; Table S6). Specifically, class 1 CRISPR-Cas systems, encompassing types I, III, and IV, were identified in 1,045 archaeal genomes spanning 14 phyla, with Halobacteriota (*n* = 527; 13 subtypes), Thermoproteota (*n* = 380; 18 subtypes), Methanobacteriota (*n* = 136; 7 subtypes), and Methanobacteriota_B (*n* = 108; 8 subtypes) exhibiting the highest abundance and subtype diversity. In contrast, class 2 CRISPR-Cas systems (types II, V) were found in only 25 genomes across four phyla, including Thermoplasmatota (*n* = 14; subtype II-A or V-A), Nanobdellota (*n* = 7; V-A or V-F1), Micrarchaeota (*n* = 2; II-C2 or V-F1) and, Halobacteriota (*n* = 2; II-A). This distribution indicates that most archaeal phyla and genomes primarily resist provirus infection by employing more complex type I systems, which function through multiple effector proteins, while only a few phyla utilize type II systems containing a single multi-domain effector protein (e.g., Cas9, Cas12a, or Cas12f1; Figure S5), whose function is equivalent to the entire effector protein complex of type I systems.

Besides CRISPR-Cas, archaea have evolved other antiviral arsenals during their arms race with viruses. A total of 35,299 non-CRISPR-Cas antiviral systems were identified across 51.5% (*n* = 11,328) of archaeal genomes, spanning 177 distinct defense system families. Among these, the restriction-modification (RM) system family is the most prevalent, present in 35.0% of genomes, followed by Prometheus (8.3%), SoFIC (6.7%), and Ceres (5.6%) (Table S7). Notably, 160 system families were detected in fewer than 1% of the genomes. Some systems, while less abundant, are widely distributed across most phyla (e.g., Shedu). In contrast, some other systems are confined to specific phyla, such as Toutatis, Erebus, and Brigantia, which are exclusively restricted to Thermoproteota. Strikingly, 256 of all identified antiviral systems in archaea are encoded by proviruses (Table S8), which not only protect the host cell but might also shield further viral infection from other proviruses^40^.

Interestingly, we found a strong positive correlation between the number of antiviral systems in a genome and the diversity of defense families in that genome (Spearman *ρ* = 0.985, *p* < 0.0001; Figure S4), indicating that genomes with more defense systems may tend to employ distinct functional families. Exceptions to this trend were also identified, with a certain number of genomes encoding numerous antiviral systems yet exhibiting limited family diversity. This type of genome is reported to typically harbor a wide diversity of RM systems^41^. Indeed, among genomes encoding more than 10 antiviral systems belonging to five or fewer families (*n* = 152), RM systems accounted for over 50% of all antiviral systems in 96.1% of cases. Further analysis revealed that both the number (Spearman *ρ* = 0.421, *p* < 0.0001) and family diversity (Spearman *ρ* = 0.416, *p* < 0.0001) of antiviral systems were positively correlated with genome size (Figure S5a and b). Conversely, no linear relationship was detected with GC content; instead, archaea with either high or low GC content contained significantly more systems and families than those with medium GC content (Figure S5c and d). Consistent with previous reports^41^, the number of proviruses per genome was positively correlated with both the number of antiviral systems (Spearman *ρ* = 0.194, *p* < 0.0001) and the number of defense families (Spearman *ρ* = 0.190, *p* < 0.0001; Figure S5e and f). Stepwise forward regression analysis confirmed that, even after controlling for genome size, the number of proviruses remained significantly correlated with both the abundance and diversity of antiviral systems (*p* < 0.0001), indicating that the antiviral arsenal in archaea is shaped by both genomic features and the number of proviruses present in the genome.

### Anti-defense systems of archaeal proviruses

Viruses can circumvent or block host defense pathways and ensure successful infection by encoding an arsenal of anti-defense proteins^42^. We detected 747 anti-defense system coding genes in 7.3% of provirus genomes (*n* = 710), which were categorized into 12 types and 39 subtypes (Figure 3a). Among these, anti-CRISPR types were the most prevalent both in abundance and diversity (288 genes, 13 subtypes), followed by anti-RM (253 genes, 9 subtypes) and anti-TA (73 genes, 4 subtypes) (Figure 3a). Most anti-defense systems were detected across a broad range of proviruses, while some were restricted to specific viral lineages. For example, anti-CRISPR, anti-RM, and anti-CBASS were commonly detected, whereas anti-RecBCD were exclusively identified in members of the family *Leisingerviridae*. Furthermore, the vast majority of proviruses (95.4%, *n* = 677) encode a single anti-defense system, while only 33 proviruses contain two or more types of anti-defense systems.

**Figure 3.**
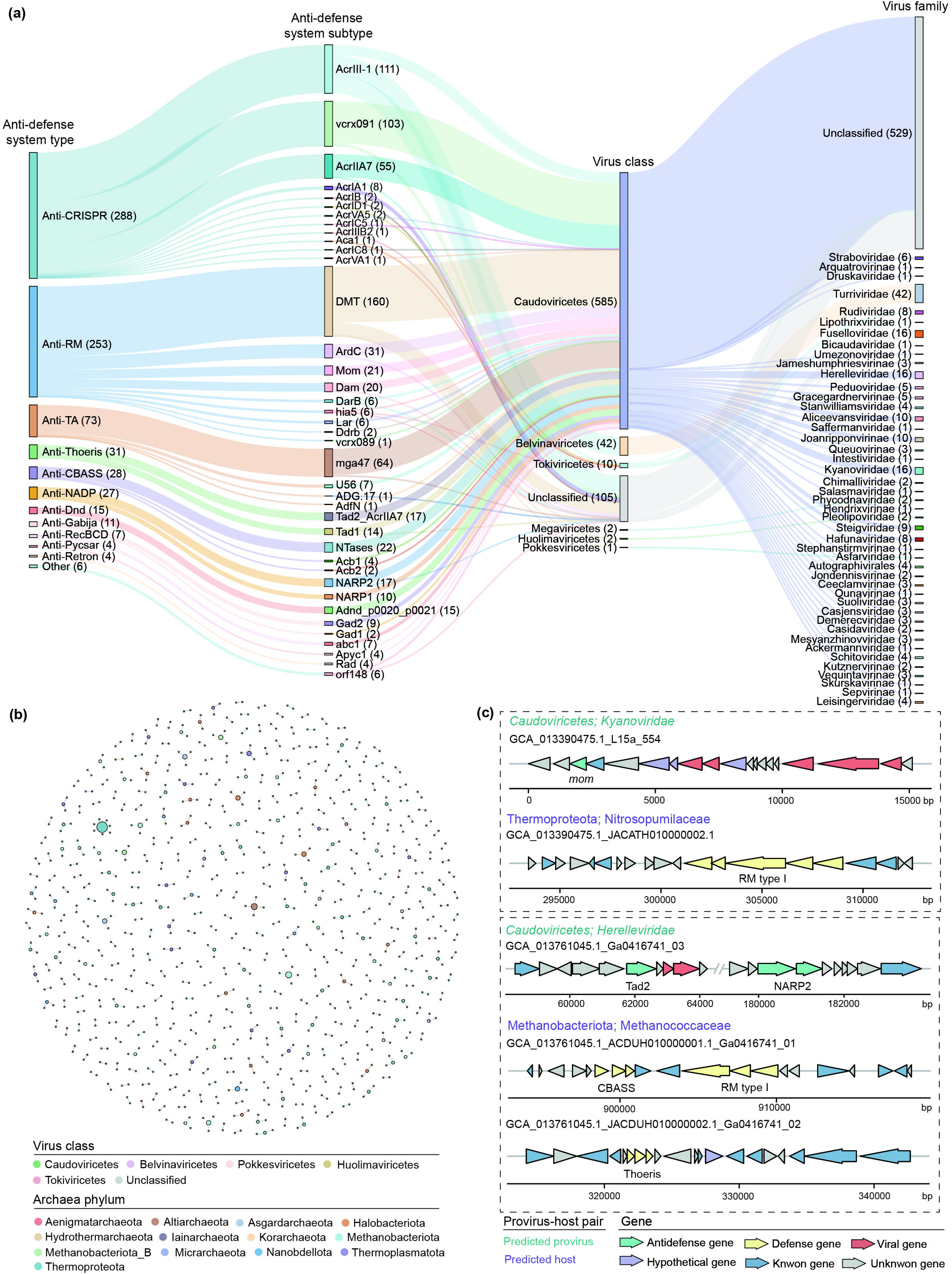
Interactions between archaeal provirus anti-defense systems and host antiviral systems. (a) Anti-defense genes identified in archaeal proviruses, showing the types of anti-defense systems and corresponding viral taxonomy. (b) A network diagram of 533 provirus-host pairs, where the provirus contains anti-defense genes and the corresponding host possesses antiviral system. The size of dots represents the strength of interactions. Different colored dots represent different viral class or archaeal phyla. (c) Genomic architecture examples of provirus gene clusters containing anti-defense systems and their corresponding host antiviral systems.

We further identified 533 provirus-host pairs in which the host encoded a defense system and the corresponding provirus carried the anti-defense system. These interactions were predominantly observed between proviruses of the class *Caudoviricetes* and their hosts, including the archaeal phyla Methanobacteriota (101 pairs), Halobacteriota (86 pairs), and Thermoplasmatota (66 pairs) (Figure 3b). Among these, only 42 proviruses possess anti-defense systems targeting their respective host’s defense machinery. For instance, a provirus from the family *Kyanoviridae* encoded a methylcarbamoylase *mom* gene predicted to counteract the RM antiviral system of its host belonging to the family *Nitrosopumilaceae*^43^. Additionally, a *Herelleviridae* provirus genome harbored NARP2 and Tad2 genes, which are inferred to target the RM and Thoeris antiviral systems of their host^44, 45^, respectively. Surprisingly, we observed that some archaeal genomes lacking identifiable antiviral systems were infected by multiple proviruses encoding anti-defense systems. One such *Methanosarcinaceae* genome (GCA_035707045.1) harbored six distinct proviruses carrying anti-defense genes, including one provirus (GCA_035707045.1_k141_17687) that carried two such genes, NARP2 and Tad2 (Figure 3c). This finding suggests that anti-defense systems may exhibit functional complexity beyond directly counteracting host immunity.

### Auxiliary Metabolic Genes (AMGs) of archaea proviruses

A total of 532 putative AMGs were identified in 321 proviruses, which potentially participate in 59 biological pathways associated with metabolism (49), environment (2), and genetic (1) information processing, organismal systems (3), and cellular processes (2) (Figure 4a; Table S9). These pathways spanned 17 functional categories, with the metabolism of carbohydrates (160, 30.0%), amino acids (96, 18.0%), membrane transport (86, 16.1%), and cofactors and vitamins (51, 9.6%) representing the most prevalent types (Figure 4b).

**Figure 4.**
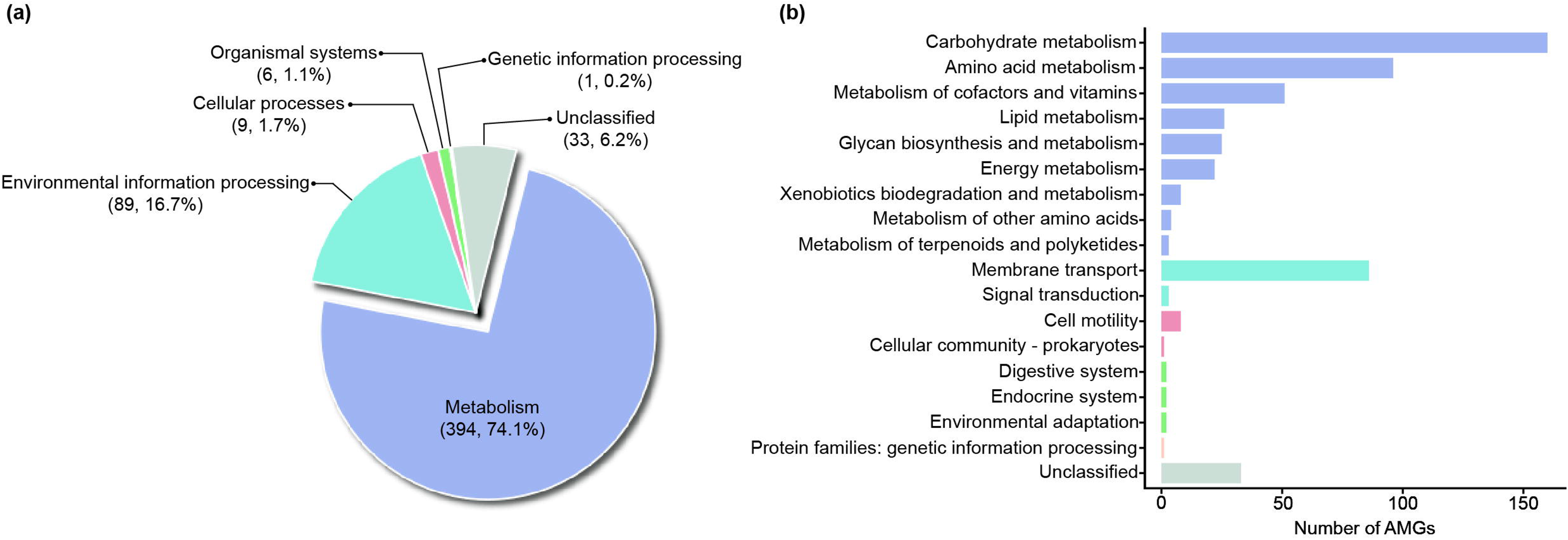
An overview of archaea provirus-encoded AMGs. (a) The percentage of viral AMGs involved in metabolism, genetic information processing, environmental information processing, cellular processes, and organismal systems. (b) Distribution of 532 viral AMGs in 17 functional categories.

Within the carbohydrate metabolism category, AMGs were predominantly enriched in central pathways, such as glycolysis/gluconeogenesis, the pentose phosphate pathway, and the metabolism of glyoxylate and dicarboxylates, fructose, and mannose. Key enzymes involved in central carbon metabolism were well-represented, including fructose-bisphosphate aldolase (FBPA) glyceraldehyde-3-phosphate dehydrogenase (GAPDH), phosphoglycerate kinase (PGK) and in glycolysis, as well as transaldolase (TAL), transketolase (TKT), and ribose 5-phosphate isomerase A (RipA) in the pentose phosphate pathway (Table S9). Importantly, this category also included AMGs encoding ferredoxin-dependent pyruvate oxidoreductase (POR) and 2-oxoglutarate oxidoreductase (OFOR), which catalyze the oxidative decarboxylation of key α-ketoacids to generate acyl-CoA and reduced ferredoxin—a critical reaction in the core carbon flow and energy conservation of anaerobic and thermophilic archaea^46–50^.

Beyond carbon metabolism, AMGs implicated in other key biogeochemical cycles were detected, including nitrogen (nitrous oxide reductase NosZ and nitrogenase iron protein NifH), sulfur (phosphoadenosine phosphosulfate reductase CysH), phosphorus (phosphate starvation-inducible protein PhoH) metabolism, and oxidative phosphorylation (NADH-quinone oxidoreductase subunit C/D NuoCD and inorganic pyrophosphatase PPA) (Table S9). The widespread presence of AMGs associated with biogeochemical cycles may enhance host nutrient uptake and utilization, thereby promoting viral replication.

We observed functional redundancy between AMGs and host metabolic genes, as revealed by comparing predicted AMG sequences with their host genomes. Specifically, 229 AMGs shared homology with host genes (> 30% amino acid identity over 50% of viral query sequence length), including 20 AMGs exhibiting > 70% sequence identity (Table S9). These highly conserved AMGs encode proteins involved in several key metabolic pathways, including the metabolism of pyruvate (phosphoenolpyruvate carboxylase PPC), glyoxylate and dicarboxylate (methylmalonyl-CoA mutase MCM), cysteine and methionine (S-adenosylmethionine synthetase MetK), and energy (NuoCD) (Table S9). The high sequence conservation of these AMGs with host metabolic genes suggests that viruses may selectively retain and exploit host-derived metabolic functions to overcome biosynthetic bottlenecks and optimize intracellular conditions for efficient viral replication^51^. Conversely, 224 AMGs showed no detectable homology to host genes, yet phylogenetic analysis revealed that some of them were consistent with the evolutionary origins of their host phyla. For instance, AMGs encoding 2-oxoglutarate ferredoxin oxidoreductase subunits (KorA and KorB) exhibit a phylogenetic affiliation consistent with the Thermoproteota, the same phylum as their hosts (Figure S6). This may be due to the loss of host genes during evolution, while viruses retained these genes and maintained their functions through horizontal gene transfer (HGT) between viruses and archaeal hosts. Notably, AMGs encoding acyl carrier protein (AcpP) and phosphate starvation-inducible protein (PhoH) not only lack homologs in their archaeal hosts but also show potential evolutionary origins from bacterial domain (Figure 5). This finding suggests that virus-mediated HGT introduces new functions to the archaeal host metabolic repertoire, facilitating ecological niche adaptation.

**Figure 5.**
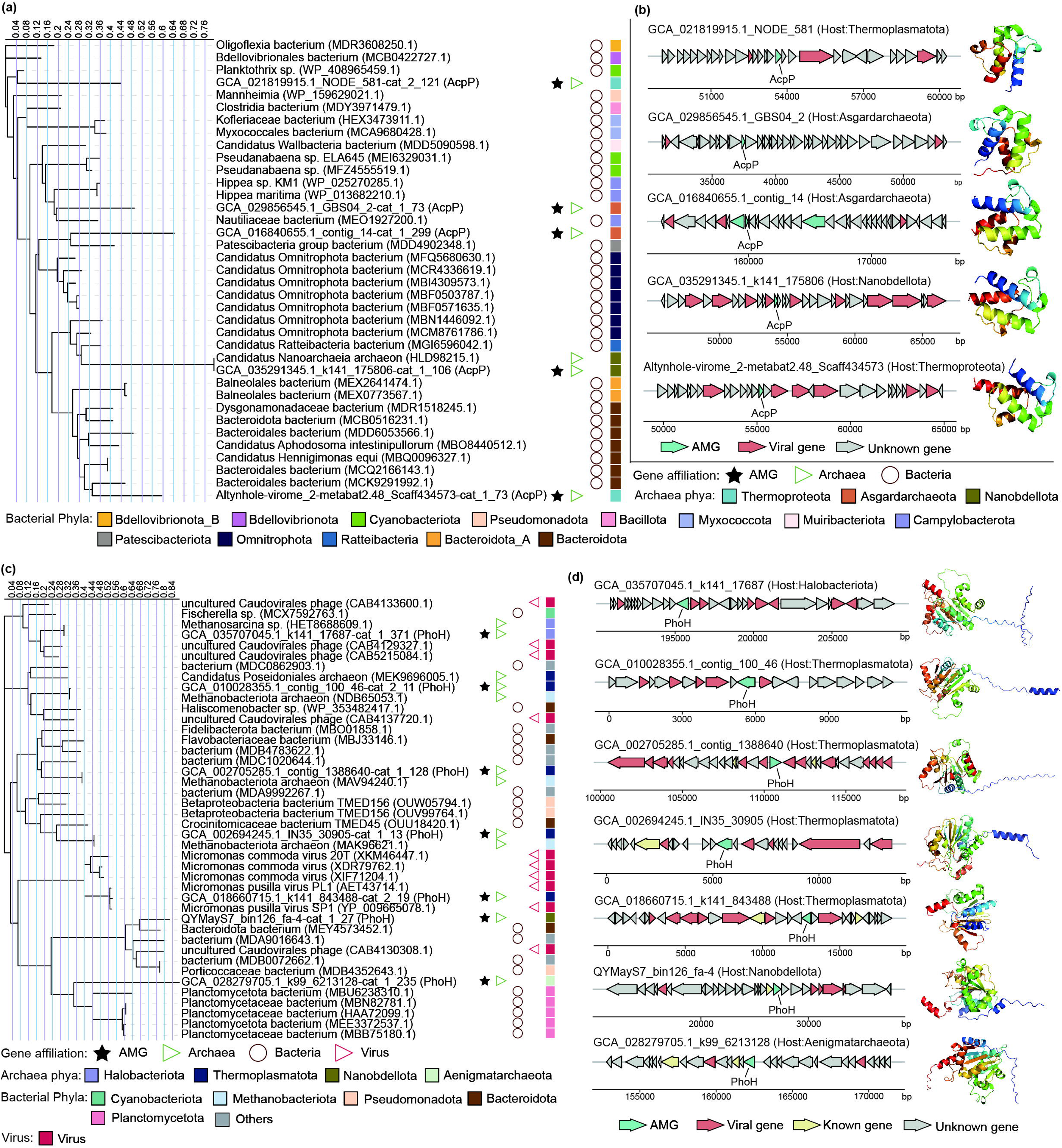
The AMGs that are without homologous to the putative host. (a) The phylogenetic tree of AcpP. (b) The genome architecture for the proviruses encoding *acpP* gene, and reference protein models for AcpP. (c) The phylogenetic tree of PhoH. (d) The genome architecture for the proviruses encoding *phoH* gene, and reference protein models for PhoH.

## CONCLUSION

In summary, this study provides a comprehensive analysis of archaeal proviruses and maps their complete genomic landscape, elucidating their taxonomic diversity, host interactions, and functional potential. We found that archaeal proviruses are not only widely distributed and highly diverse but also harbor a large number of unexplored viral lineages. Host range analysis revealed that many proviruses exhibit broad infectivity across archaeal lineages, with some even capable of cross-domain infection. The discovery of antiviral and anti-defenses systems underscores the ongoing evolutionary arms race between archaeal hosts and their proviruses. In addition, auxiliary metabolic genes were widely detected and found to be involved in key metabolic processes such as carbon, nitrogen, and sulfur metabolism, suggesting that proviruses may indirectly modulate biogeochemical cycling through metabolic reprogramming of their hosts. These findings significantly advance our understanding of archaeal provirus diversity and provide a foundational resource for further exploring virus–host dynamics and the ecological impacts of Archaea.

## METHODS

### Archaea genome collection

This study collected a total of 25,530 archaeal genomes, comprising 24,840 genomes (including cultured isolates, metagenome-assembled genomes (MAGs), and single-amplified genomes (SAGs)) obtained from the NCBI GenBank database (November 2024), and 690 archaeal MAGs from our previous publications^52, 53^. Following data collection, we performed quality screening and taxonomic validation on all archaeal genomes. Specifically, CheckM v1.0.12^54^ was used to evaluate genome completeness and contamination, and low-quality genomes (completeness < 50% or contamination > 10%) were excluded. The retained genomes were subsequently dereplicated using dRep v3.3.0^55^ with an average nucleotide identity (ANI) cutoff value of 99.99%. Taxonomic classification and validation were performed using GTDB-Tk v2.4.1 (reference data r226)^8^, confirming all as Archaea. Ultimately, our study generated a non-redundant set of 21,989 high-quality archaeal genomes (21,593 from the NCBI GenBank database and 396 from our laboratory). These archaea are classified into 21 phyla, 65 classes, 172 orders, and 603 families. Among these, there were 48 families with an abundance of over 100 available genomes, 33 families with 50 to 100 genomes, 25 families with 30 to 50 genomes, and the remaining 496 families with fewer than 30 genomes (see Table S1 for details).

### Identification, quality checking and clustering of proviruses

Provirus regions (≥ 5 kb) were first identified from archaeal genomes using geNomad v1.11.0^56^ and Virsorter2 v2.2.4^57^ with default parameters. Putative provirus sequences from the two pipelines were merged, and duplicate sequences were removed via an all-versus-all BLASTn analysis (parameters: -perc_identity 100 -qcov_hsp_perc 100 -evalue 1e-3). The resulting sequences were then processed with CheckV v1.0.1 (database v1.5)^58^ for quality assessment, during which only proviruses containing viral hallmark genes were retained. The provirus genomes were further screened and retained based on the following criteria: (i) geNomad virus score ≥ 0.8 and to either encode one virus hallmark (for example, terminase, capsid proteins, portal protein and so on), or geNomad viral marker score ≥ 5.0, (ii) score > 0.9 for VirSorter2, or score between 0.7–0.9 with at least one viral hallmark gene, (iii) high- or medium-quality CheckV completeness estimates or the presence of direct terminal repeats (DTRs). The obtained proviruses were clustered at 95% ANI and 85% alignment fraction (AF) of the shortest proviruses using the CheckV script (https://bitbucket.org/berkeleylab/checkv/src/master/) to generate non-redundant viral operational taxonomic units (vOTUs).

### Lifestyle prediction of proviruses

The lifestyle of proviruses was predicted using four methods. (i) CheckV v1.0.1 predicts lysogenic lifestyle by detecting provirus boundaries. (ii) The proviruses with proteins annotated (see below, DRAMv annotation of viral proteins) as integrases, invertase, serine recombinases, transposase, *parA/B*, and CI/Cro repressor are considered proviruses^59^. (iii) PhaTYP v0.3.0 (minimum score ≥ 0.9) and VIBRANT v1.2.1 with default parameters were used to infer the lifestyle of the provirus genome. The combined result of more than two software was used as the final prediction result. Conflicting results between two software were resolved through sequential priority assignment to CheckV, gene annotation, VIBRANT, and PhaTYP outputs.

### Taxonomic classification of vOTUs

The obtained vOTUs were taxonomically classified using geNomad v1.11.0^56^, vConTACT2 v0.11.3^60^, VITAP v1.7.1^61^, PhaGCN3 v3.1, PhaGCN2 v2.3^62^, and viral genome homolog search method based on BLASTn comparison of the IMG/VR V4 database^36^. Briefly, geNomad with the default parameter was used to classify vOTU by taxonomic rank of annotated proteins. For vConTACT2, open reading frames (ORFs) were first predicted with Prodigal (v2.6.3; -p meta)^63^, and the resulting protein sequences were used for vConTACT2 and the NCBI “ProkaryoticViralRefSeq201-Merged” was selected as the reference database. A vOTU was assigned to a viral family if over 50% of its proteins matched proteins from that family with a bit score ≥ 50^59^. VITAP was employed with default parameters, applying a lineage score cutoff of ≥ 0.9 for taxonomic classification. PhaGCN3 and PhaGCN2 classified vOTUs at the family-level using recommended cutoff scores >0.5^62^. For the BLASTn-based approach, taxonomy was assigned based on the top best hit in the IMG/VR V4 database that met the thresholds of ≥ 90% ANI and ≥ 75% AF^64^. The final taxonomic assignment for each vOTU was determined through hierarchical integration of results from the six pipelines, prioritized in the following order: vConTACT2, BLASTn, geNomad, VITAP, PhaGCN3, and PhaGCN2. If conflicts existed at a lower taxonomic level (e.g., family level) for different pipelines but there was a common higher level of taxonomy (e.g., order level), then the higher taxonomic level was assigned.

### Comparison of provirus genomes and proteins to public databases

Archaeal provirus sequences and their encoded proteins were compared against the existing archaeal virus genomes and protein databases in IMG/VR v4^36^ and NCBI Viral RefSeq. First, all known archaeal virus sequences from these two databases were clustered into vOTUs using the above method with 95% ANI and 85% AF. Next, the 9,123 vOTUs identified in our study were clustered against this reference set of vOTUs. Finally, the amino acid sequences of our archaeal vOTUs were annotated with Prodigal v2.6.3^63^ and subsequently clustered alongside the sequences from IMG/VR v4 and NCBI RefSeq using CD-HIT v4.8.1 (parameters: -c 0.6 -G 0 -aS 0.8 -n 4)^65^.

### Identification of antiviral systems and provirus–host predictions

CRISPR-Cas genes and arrays were extracted from archaeal genomes using CRISPRcasFinder v4.3.2. Those with evidence level 4 (repeat conservation index > 70% and spacer identity < 8%) were retained for subsequent analysis. CRISPR–Cas systems in the contigs with CRISPR arrays were annotated by CRISPRCasTyper v1.8.0. Other antiviral defence systems in archaeal were detected using DefenseFinder v2.0.0 (parameters: --db-type gembase)^41, 66^, updated with all known anti-phage systems in February 2025 (defense-finder-models v2.0.2).

To predict host–provirus interactions, we first performed BLASTn alignments between all proviruses and archaeal CRISPR-spacer dataset. If a spacer generated an exact match (identity = 100%, coverage = 100%) with a provirus, we assigned the host associated with the spacer to the provirus. Furthermore, potential hosts for archaeal proviruses were also predicted using the integrated CRISPR method within iPHoP v1.4.1 (database: iPHoP_db_Jun25_rw)^67^.

### Identification of anti-defense systems

The identification of anti-defense genes in proviruses was conducted using two approaches. (i) Anti-defense genes in the provirus were predicted using DefenseFinder (--antidefensefinder-only)^41,66^. (ii) Anti-Prokaryotic Immune System (APIS) was selected as a reference database^68^, which contains experimentally validated viral proteins that counteract prokaryotic immune systems and their associated protein families. DIAMOND blastp v0.9.14 (parameters: --evalue 1e-10 --id 30 --max-target-seqs 1)^69^ and HMMER v3.4 (parameters: --domtblout --noali -E 1e-10)^70^ were used to annotate the anti-defense genes from provirus sequences following the protocol provided by dbAPIS (https://bcb.unl.edu/dbAPIS/downloads/readme.txt).

### Auxiliary metabolic genes (AMGs) identification and novelty assessment

Function annotations of gene and predicted AMGs for all proviruses using the DRAM-v pipeline^71^. Briefly, all proviruses were re-analyzed using VirSorter2 (parameters: -prep-for-dramv - min-length 5000 -min-score 0 -include-groups dsDNAphage,ssDNA,NCLDV,RNA,lavidaviridae) to generate an “affi-contigs.tab” file, which was subsequently annotated using DRAM-v with the default database. The “-min-score 0” parameter was employed to ensure maximal sensitivity, as some proviruses derived from geNomad were not classified as viral contigs by VirSorter2 when using the “-min-score 0.5” parameter, as previously described^72^. Predicted AMGs were filtered based on the following criteria: auxiliary_score ≤ 3 and amg_flag assigned to M or M with E and/or K. Additionally, the flanking genes of the putative AMG were manually inspected to ensure the presence of viral marker genes or viral-like genes. To enhance the confidence of AMG identification, we further performed manual curation to exclude genes potentially involved in essential viral processes, including viral invasion (i.e., glycoside hydrolases and peptidases involved in cell wall lysis), modification of viral components (i.e., glycosyl transferases, adenylyltransferases and methyltransferases that putatively involved in viral DNA, RNA, and structural proteins modification), viral replication, transcription, and translation (e.g., nucleotide metabolism, and ribosomal proteins), and folate biosynthesis^27, 73, 74^.

To provide insight into the potential functional relevance of AMG to the host, we first compared provirus sequences to the host genome using BLASTn (parameters: -perc_identity 100 - qcov_hsp_perc 100 -evalue 1e-3) to determine that the host genome was unintegrated or uncontaminated with viral sequences. Subsequently, the AMG was compared to the host genome using BLASTp to assess the relatedness of the identified AMG to homologous genes found in the host^51^. The three-dimensional structures of AMGs were predicted using AlphaFold3^75^.

### Phylogenetic analysis of AMGs

To investigate the probable evolutionary origin of the AMGs, homolog sequences of AMGs were recruited from the NCBI nr database (Release October, 2025), and the top 10 hits with a bit score > 50 were retained as reference sequences for the AMGs. The protein structures for all retrieved sequences were predicted using AlphaFold3. All AlphaFold structures were fed to Foldtree^76^ to construct a phylogenetic tree using the parameters filter=False, align_mode=1, and custom_structs=True. Briefly, Foldseek compresses the local structural of each residue into a structural alphabet (3Di) via a pre-trained vector quantized variational autoencoder (VQ-VAE)^77^, enabling fast alignment of the new structure-aware sequences. It combines amino acid (AA) and 3Di substitution scores with the default weighting scheme to rank the potential alignments (AA+3Di), with two more alignment results using LDDT and TM score. The resulting alignment outputs are used to generate distance matrices, from which phylogenetic trees are constructed via neighbor-joining with QuickTree^78^ and rooted using the minimal ancestor deviation (MAD) method^79^. Phylogenetic trees were visualized using the iTOL online server^80^. We regard the sources of sequences that cluster closely with AMG in the phylogenetic tree as potential sources of AMG.

### Statistical analyses and visualization

All statistical analyses in this study were performed using R v4.1.4. Correlations between host GC content and provirus-related traits—including provirus number, provirus proportion, and antiviral system and family counts—were evaluated using Mann–Whitney U tests via the wilcox.test function from the R “stats” package. Additionally, Spearman correlation analyses were carried out using the cor.test function (“stats” package) to evaluate (i) the relationships between host genome size and the same set of provirus-related traits, and (ii) the correlations between provirus number and the counts of antiviral systems and family. To assess provirus enrichment across different host taxonomic levels (phylum, class, order, family, genus), we applied a bootstrapping-enhanced Mann–Whitney U test, restricting the analysis to genera with at least 50 genomes. In brief, for the bootstrapping, we collected 10 random genomes for each genus at each iteration. This step was repeated 100 times (bootstrapping). To identify determinants of antiviral system abundance while controlling for genome size, we conducted stepwise forward negative-binomial regression using the step function in the R “MASS” package, retaining variables selected by AIC criterion. The manuscript figures were mainly generated using the R packages “ggplot2”, “ggpubr”, “dplyr”, “igraph” and “gggene”. The figure format was adjusted using Adobe Illustrator if needed.

## ACKNOWLEDGMENTS

This study was supported by the National Natural Science Foundation of China (42421001, 42222105, 42171144) and the Global Ocean Negative Carbon Emissions (Global ONCE) Program. We would like to express our gratitude to the Supercomputing Center of Lanzhou University for their valuable support in the computation works.

## AUTHOR CONTRIBUTIONS

Yang Zhao: Data curation (lead); formal analysis (lead); software (lead); conceptualization (equal); writing-original draft (lead); writing-review and editing (lead). Pengfei Liu: Funding acquisition (lead); project administration (lead); resources (lead); conceptualization (equal); writing-review and editing (equal). Meiling Feng: Data curation (supporting) and writing-review and editing (supporting). Rong Wen: Data curation (supporting) and writing-review (supporting). Zhihao Zhang: Data curation (supporting) and writing-review (supporting). Xingyu Huang: Data curation (supporting) and writing-review (supporting). Hongfei Chi: Data curation (supporting) and writing-review (supporting).

## DATA AVAILABILITY

The present study did not generate codes, and mentioned tools used for the data analysis were applied with default parameters unless specified otherwise.

