## Supplementary Figures for "Unraveling the archaeal virosphere: diversity, functional and virus-host interactions"

Zhao et al.

### Supplementary figures

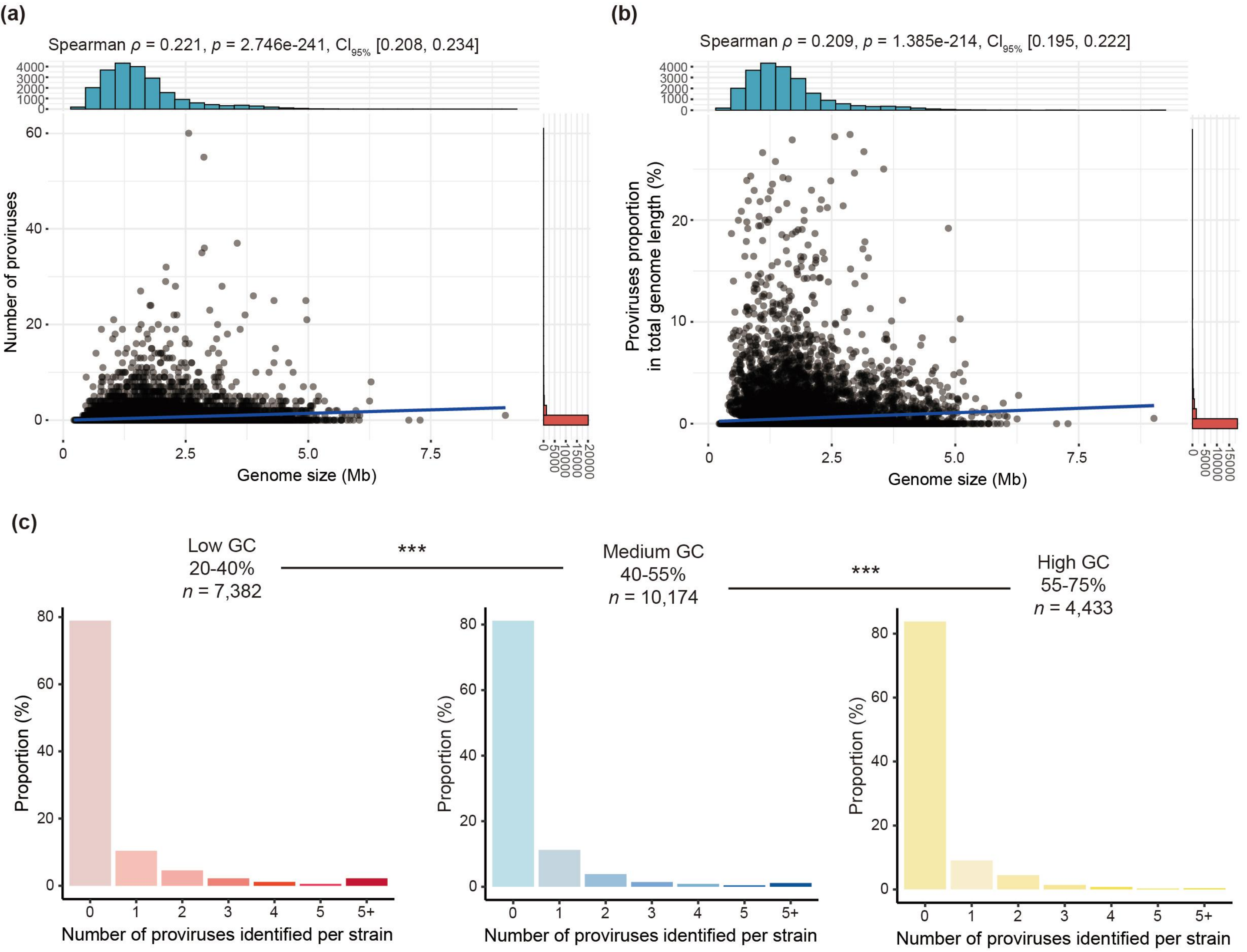


**Figure S1 |The overall landscape of archaeal proviruses (related to Figure 1).** (a) The correlation between genome size and the number of proviruses in that genome (Spearman's correlation coefficient $\rho = 0.221$, $p < 0.0001$). (b) The correlation between genome size and the proportion of proviruses in the genome (Spearman's correlation coefficient $\rho = 0.209$, $p < 0.0001$). Each dot represents a strain. (c) Comparison of the number of proviruses carried by the "Low GC", "Medium GC", and "High GC" archaea. Statistical significance tests were performed using the nonparametric Mann-Whitney U test, and the two-tailed $p$ values were calculated. ***: $p < 0.001$.

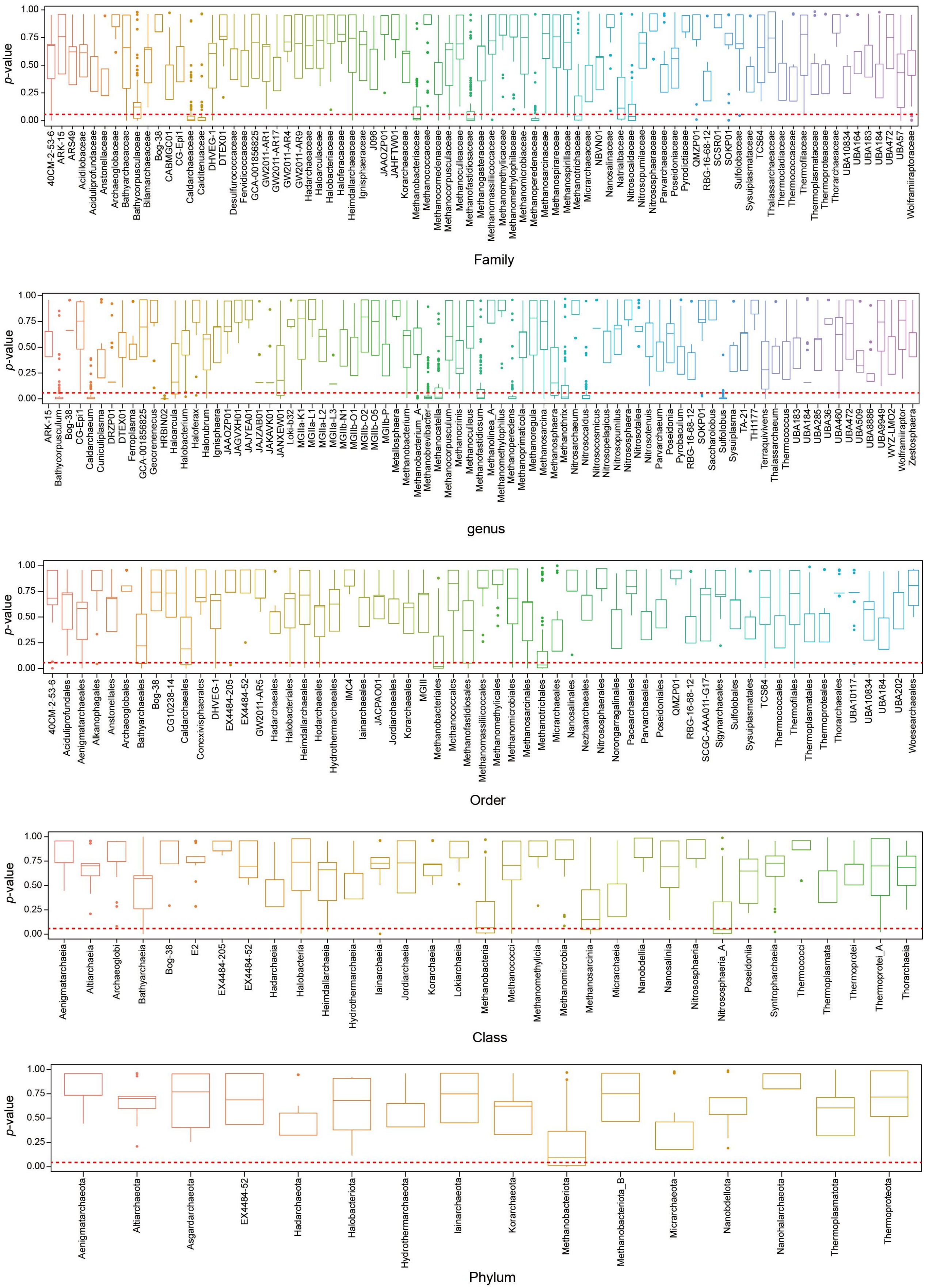


**Figure S2 | Boxplots of *p*-values at different classification levels.** *p*-values were calculated collecting and bootstrapping (100 times randomly) the number of proviruses in 10 genomes for each classification level, using the Mann-Whitney U test (see Methods section). The red dotted line indicates a significant Mann-Whitney U test ($p < 0.05$).

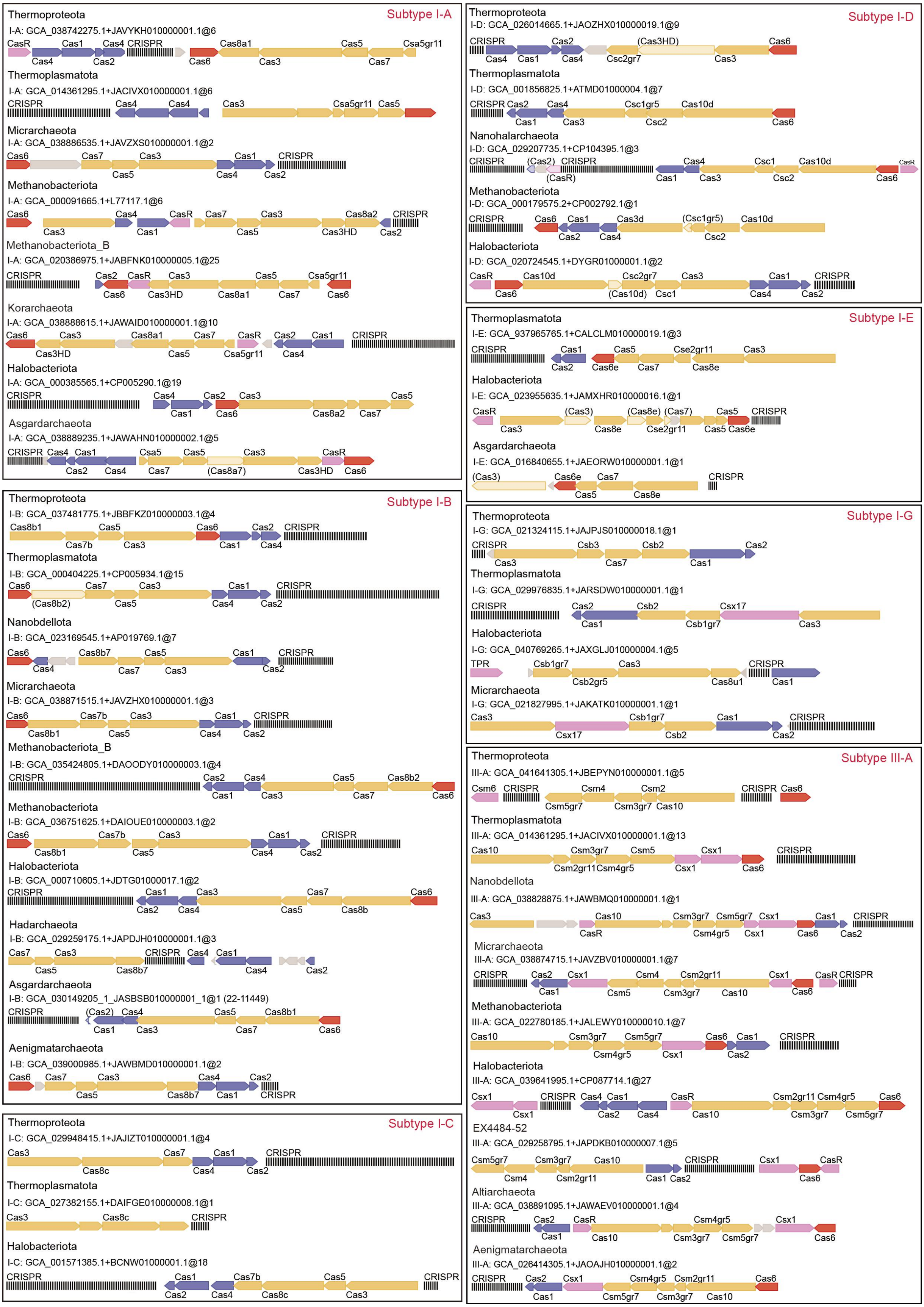


**Figure S3 | Genetic architectures of CRISPR–Cas systems in archaea.** Genetic architectures of each subtype of the identifiable CRISPR–Cas system in archaeal phyla. If one subtype is possessed by multiple genomes for an individual phylum, the representative one is shown. CRISPR–Cas system subtypes belonging to classes 1 and 2 are in red and blue text, respectively, on the top right corner within each frame.

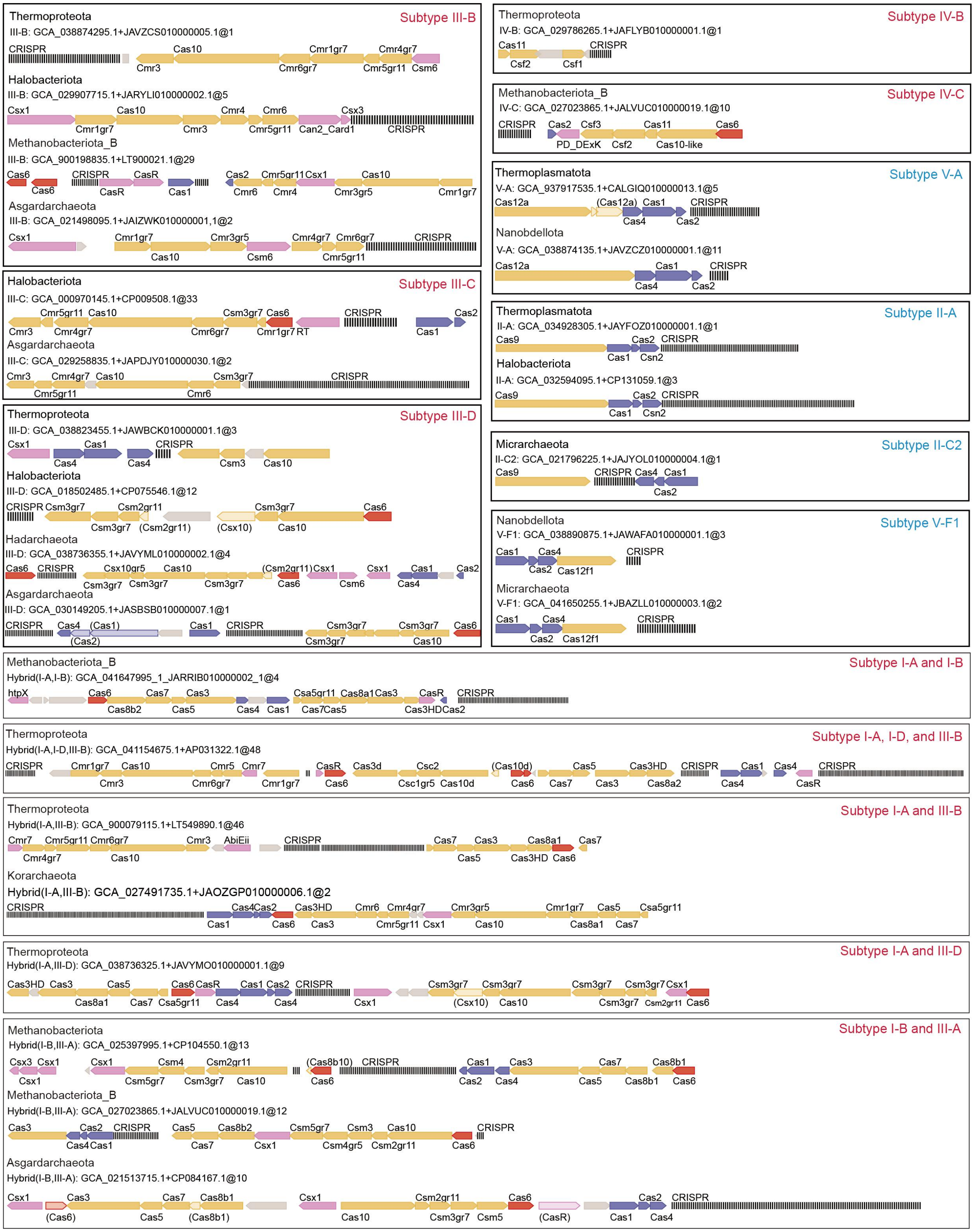


**Figure S3 | Continue.**

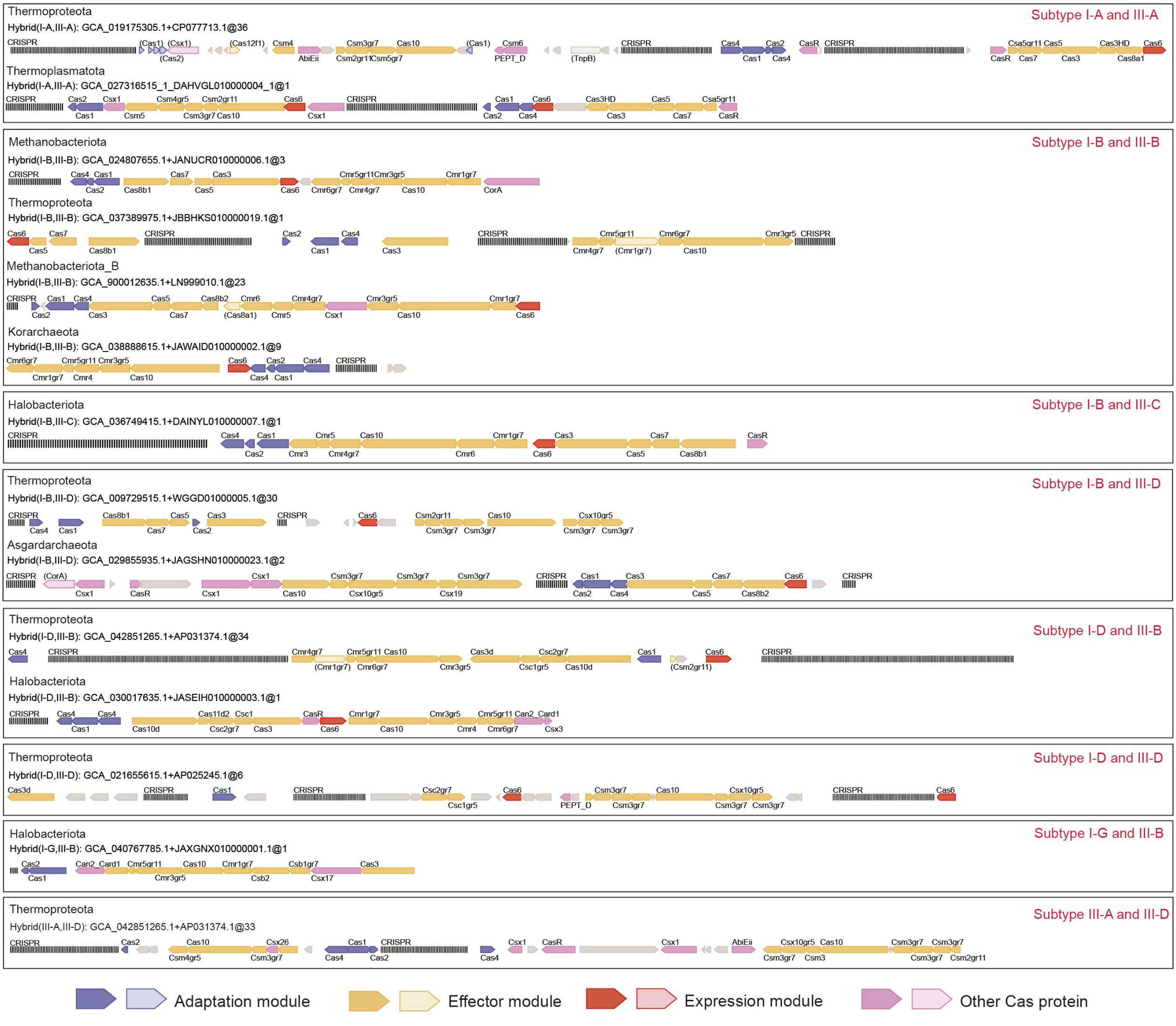


**Figure S3 | Continue.**

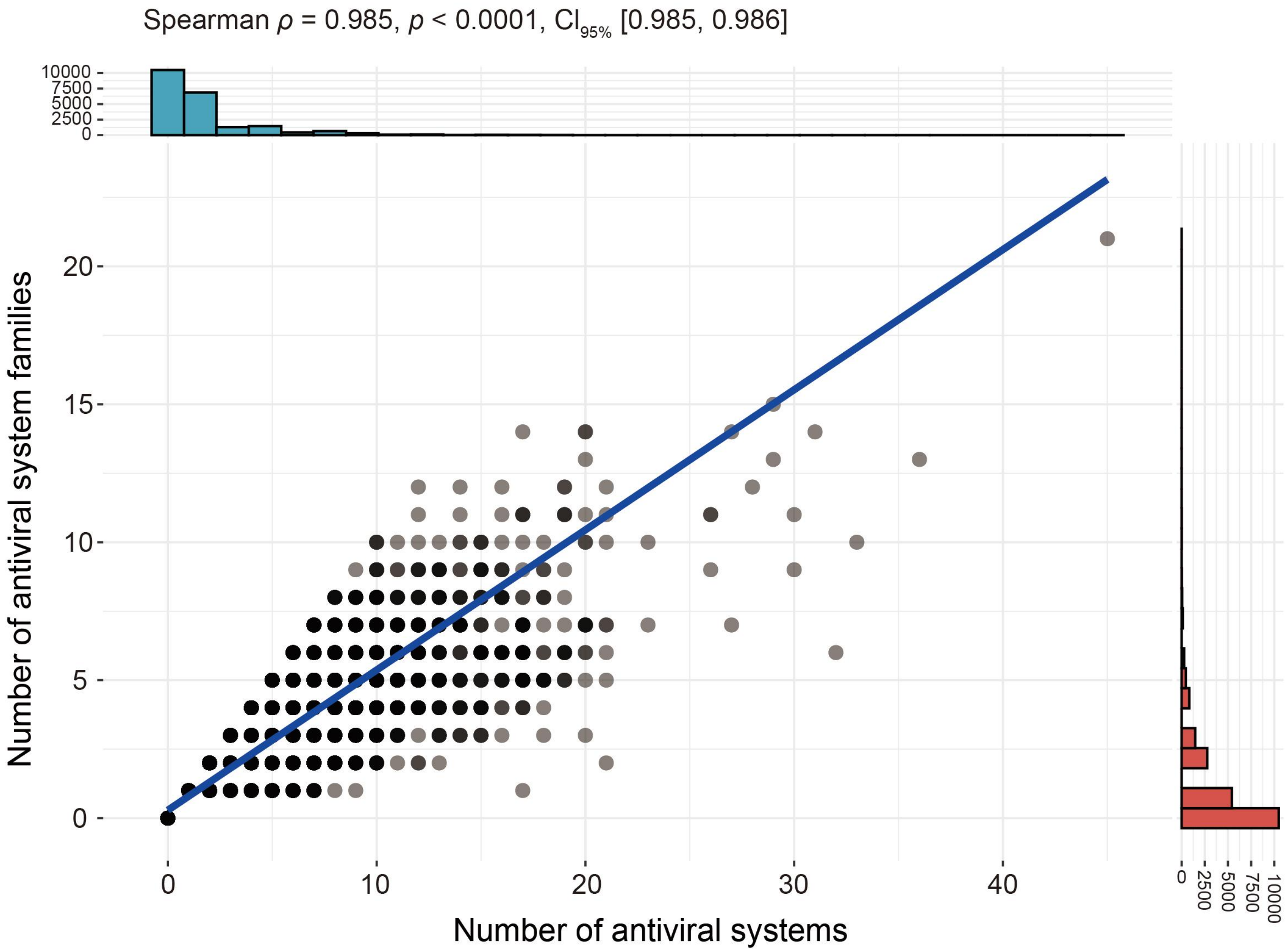


**Figure S4 | Correlation between the families of antiviral systems and the total number of antiviral systems (Spearman $\rho = 0.985$, $p < 0.0001$). Each dot represents a strain.**

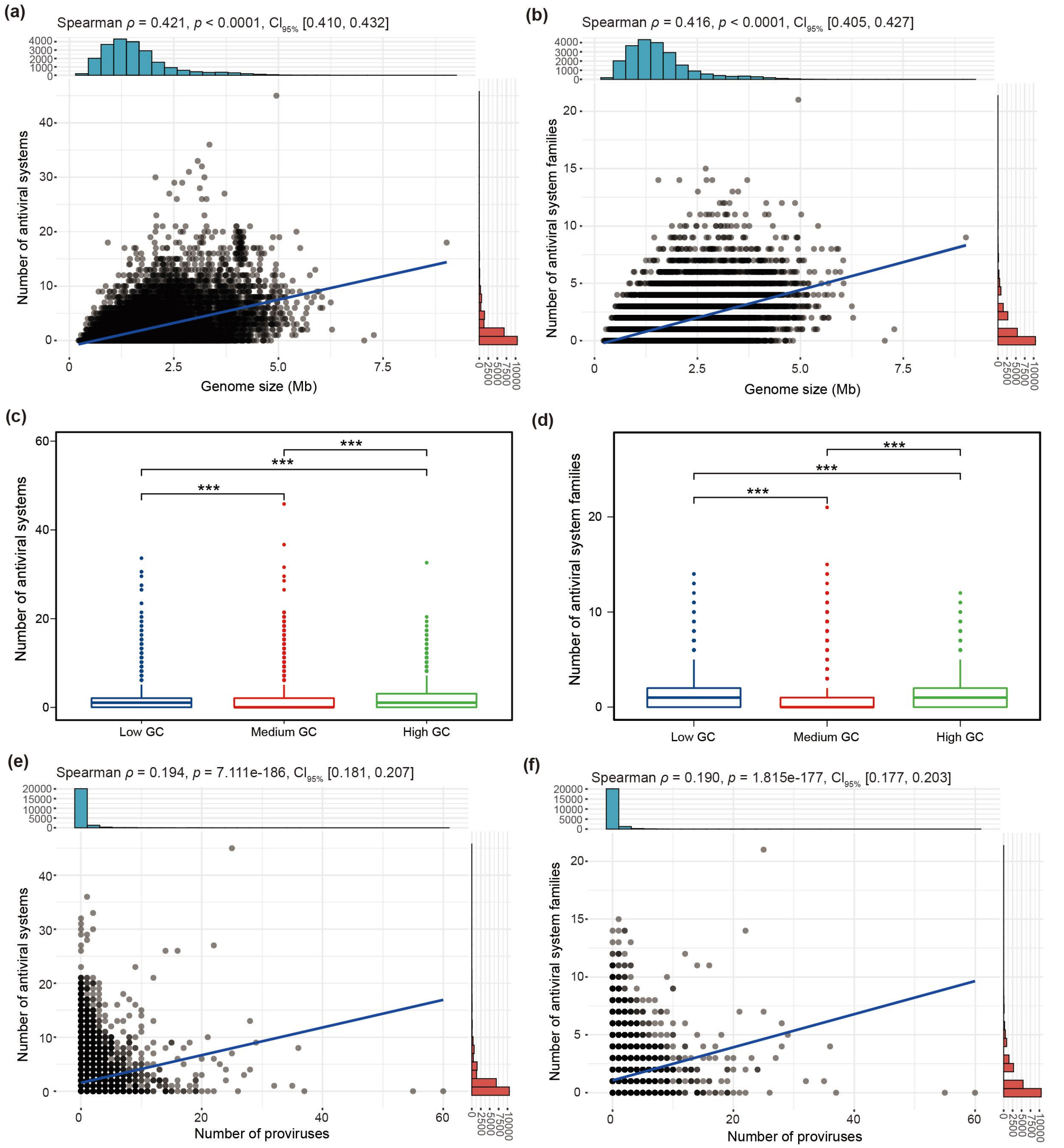


**Figure S5 | Potential drivers of antiviral system distribution.** (a) The correlation between genome size and the number of antiviral systems in the genome (Spearman $\rho = 0.421$, $p < 0.0001$). (b) The correlation between genome size and the number of antiviral system families in the genome (Spearman $\rho = 0.416$, $p < 0.0001$). (c) Comparison of the number of antiviral systems carried by the "Low GC", "Medium GC", and "High GC" archaea. Statistical significance tests were performed using the nonparametric Mann-Whitney U test, and the two-tailed $p$ values were calculated. ***: $p < 0.001$. (d) Comparison of the number of antiviral system families carried by the "Low GC", "Medium GC", and "High GC" archaea. Statistical significance tests were performed using the nonparametric Mann-Whitney U test, and the two-tailed $p$ values were calculated. ***: $p < 0.001$. (e) The correlation between the number of proviruses in the genome and the number of antiviral systems (Spearman $\rho = 0.194$, $p < 0.0001$). (f) The correlation between the number of proviruses in the genome and the number of antiviral system families (Spearman $\rho = 0.190$, $p < 0.0001$). Each dot represents a strain.

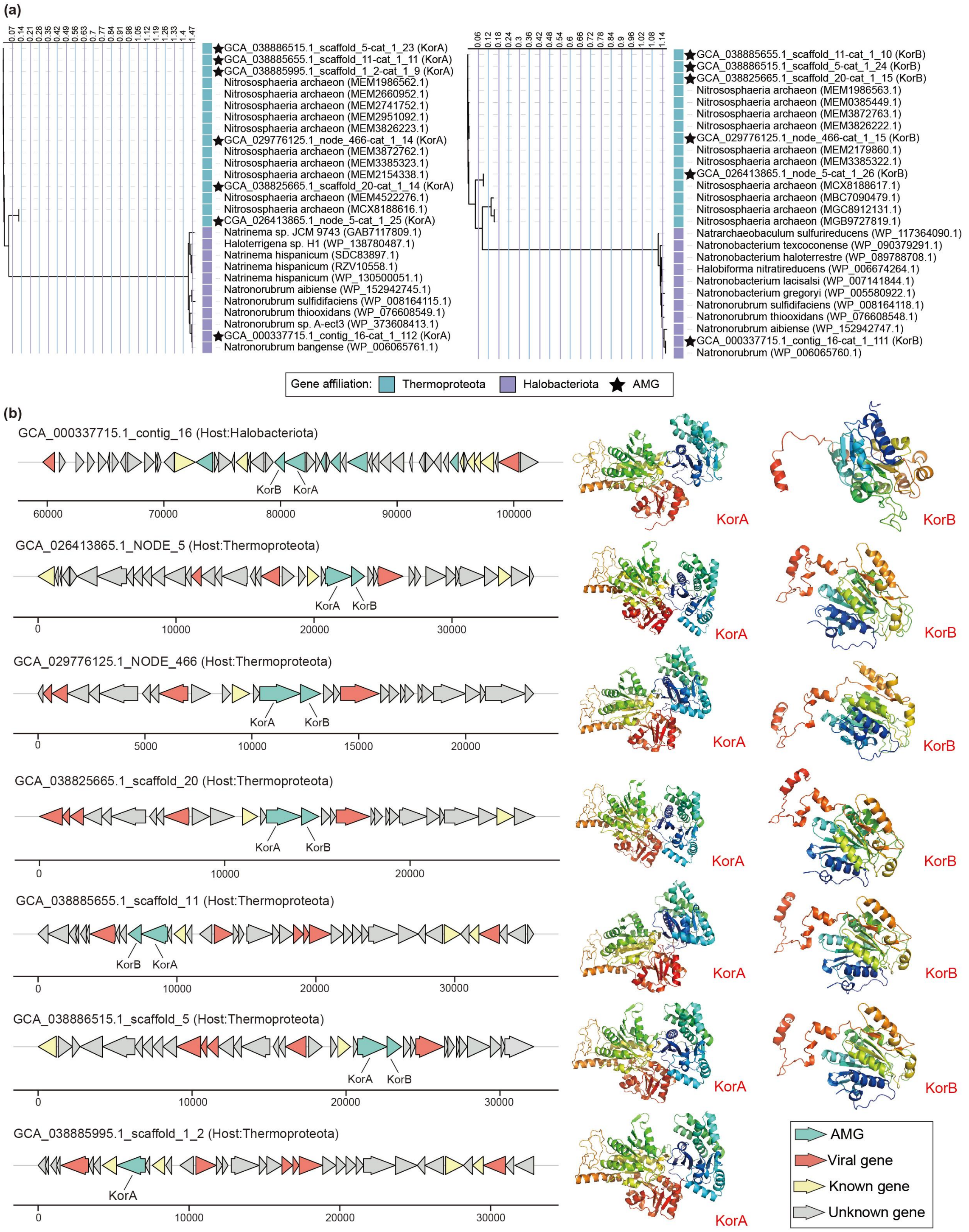


**Figure S6 | The AMGs that are without homologous to the putative host.**

(A) The phylogenetic tree of KorA and KorB. (B) The genome architecture for the proviruses encoding *korA* and *korB* genes, and reference protein models for KorA and KorB.
